# Cold Atmospheric Plasma Selectively Disrupts Breast Cancer Growth in a Bioprinted 3D Tumor Microenvironment Model

**DOI:** 10.1101/2024.12.30.629225

**Authors:** Laura M. Bouret, Jean-Baptiste Billeau, Michael H. Weber, Derek H. Rosenzweig, Stephan Reuter

## Abstract

Spine metastases are the most common bone site for breast cancer, with evolving surgery and multidisciplinary care improving outcomes. Current treatments, including chemotherapy and invasive surgery, may damage healthy tissue and may leave residual tumors that lead to recurrence. Cold atmospheric plasma (CAP) offers a non-invasive alternative by delivering reactive oxygen and nitrogen species (RONS) locally to tumor sites, selectively targeting cancer cells while sparing healthy tissue. To assess the impact and selectivity toward tumor cells adjacent to bone-like tissue, we develop a 3D bioprinted tumor-stroma model using a 1% alginate and 7% gelatin cell-laden hydrogel to mimic a bone-like microenvironment. The model co-cultures triple-negative MDA-MB-231 human breast cancer cells with primary human bone marrow mesenchymal stromal cells to simulate tumor-stroma interactions. The effects of CAP treatments are assessed through metabolic activity and viability assays over three days. Results show significant selectivity for cancer cells in both 2D and 3D cultures. CAP minimizes damage to healthy cells, offering the potential for localized treatment over systemic chemotherapies such as doxorubicin. Our novel bioprinted model, combined with a plasma source controlling RONS composition, enables detailed studies of redox-based cancer cell inactivation and highlights CAP as a personalized, non-invasive treatment for bone metastases.

## 1. Introduction

Bone, particularly the spine, is a common site of metastasis for cancers such as breast, lung, and prostate.[1,2] These metastases may be osteolytic in nature, and they cause significant bone loss, leading to skeletal-related events (SREs), including fractures, pain, hypercalcemia, and spinal cord compression – all of which severely impact patients’ quality of life and present substantial challenges for treatment.[3–5] Current therapeutic approaches include surgery, radiation, and systemic therapies such as chemotherapy, immunotherapy, and hormonal therapy.[6,7] However, these strategies face limitations. Surgical removal of metastatic lesions often necessitates the excision of healthy tissue to ensure all malignant cells are removed, resulting in large defects that require reconstruction.[8] Alternatively, a partial resection of a metastasis will result in a residual tumor, which may cause symptomatic local recurrence.

Systemic chemotherapy may be used to treat residual tumors or other pre-clinically significant sites; however, it often lacks specificity, damages healthy tissues, and can cause severe side effects.[6] Targeted antibody treatments hold promise for improving specificity and reducing side effects, but their success is often limited by eventual therapy resistance. The frequent relapses and resistance to existing therapies underscore the need for innovative approaches that are more targeted and versatile.[4,6,9] Cold atmospheric plasma (CAP) has emerged as one such approach. By leveraging its selective cytotoxic effects on cancer cells while reducing damage to healthy tissues, CAP could offer minimally invasive treatment of primary solid tumors. Also, it could intra-operatively complement surgical interventions by minimizing the extent of healthy tissue excision, preserving critical structures, and treating any residual disease left in the tumor bed, making it a valuable addition to existing treatment options for bone metastases.[10,11]

CAP is an innovative therapeutic modality created by electrically ionizing a gas, generating reactive oxygen and nitrogen species (RONS) at tissue-tolerable temperatures.[12–17] These RONS include hydrogen peroxide (H2O2), ozone (O3), singlet delta oxygen (^1^O2), atomic oxygen (O), hydroxyl radicals (^•^OH), and nitric oxide (^•^NO). For better readability, the dot notation is omitted in the remainder of the manuscript. RONS play a critical role in CAP treatment’s anticancer effects due to cancer cells’ heightened sensitivity to oxidative stress caused by altered metabolism.[18–21] CAP can be applied directly to tissues or indirectly via plasma-treated media, offering flexibility and minimizing off-target effects.[12,22–24] However, to maximize therapeutic effectiveness, CAP treatments require careful optimization of device design, gas composition, and application methods.[25–28]

While the selective cytotoxicity of CAP has been demonstrated in 2D cell cultures, translating these findings to more physiologically relevant 3D models remains challenging. Traditional *in vitro* models, such as spheroids and organoids, offer partial improvements over 2D cultures but often fail to fully replicate the complexity of the tumor microenvironments, such as matrix proteins and various diverse cell types.[29–31] Spheroids can develop hypoxic cores, a feature shared with solid tumors, which can be advantageous for studying treatment resistance but may also alter cellular behavior in context-dependent ways.[29–31] Organoids, despite their structural complexity, lack scalability and precise control over spatial organization.[29] Microfluidic tumor-on-a-chip systems simulate complex interactions more effectively but are costly and technically demanding.[29] Similarly, while animal models are essential for preclinical research due to their ability to capture whole-organism complexity, their inherent biological complexity makes it challenging to study specific mechanisms, such as cell-cell interactions or protein functions.[32] Additionally, their limited predictive value for human outcomes frequently poses translational challenges in preclinical research.[32] These limitations highlight the need for advanced 3D models, such as bioprinted systems, that balance human physiological relevance with scalability and reproducibility while complementing existing preclinical approaches.[29,32]

3D bioprinting offers a transformative solution to these challenges by enabling the fabrication of complex tissue constructs that mimic the tumor microenvironment.[33,34] This technology integrates living cells and biomaterials in defined spatial arrangements, supporting the inclusion of multiple cell types and extracellular matrix (ECM) components to simulate native tissue architecture. By incorporating patient-derived cells, 3D bioprinted models enable personalized studies of therapeutic responses, addressing patient heterogeneity and advancing precision medicine. This technology allows for integrating vascular networks and creating reproducible, scalable models critical for studying complex processes such as metastasis and treatment responses.[29,33,34] These advantages make bioprinting an ideal platform for preclinical research evaluating emerging therapies such as CAP therapy, offering unique insights into plasma-tissue interactions in a controlled, reproducible, physiologically relevant setting.[33,35]

In this study, we introduce a novel bioprinted tumor-stroma platform to investigate the selective effects of CAP therapy in a bone-mimicking microenvironment. This platform incorporates a co-culture of MDA-MB-231 human triple-negative breast cancer cells and human bone marrow mesenchymal stromal cells (hbmMSCs) within a hydrogel designed to simulate the cellular and mechanical properties of the metastatic bone marrow niche.[1,4,36]

MDA-MB-231 is an aggressive breast cancer cell line with limited treatment options and a high inclination for bone metastases, particularly in the spine. Their resistance to conventional therapies makes them particularly suited for evaluating the selectivity of CAP treatment in this context. By comparing CAP treatment effects on this 3D co-culture model with those in 2D cultures and 3D monocultures, we aim to evaluate CAP treatment’s selective action against cancer cells and its impact on tumor-stroma dynamics. Furthermore, we explore how the controlled, reproducible nature of 3D bioprinted constructs enhances the study of plasma-tissue interactions, providing insights into optimizing CAP parameters for effective treatment. This work represents the first-of-its-kind application of CAP therapy on a 3D bioprinted tumor-stroma model. It offers a foundation for advancing CAP as a localized, non-invasive treatment for bone metastases and paves the way for personalized cancer therapies. In future studies, this platform has the potential to incorporate patient-specific variability using primary bone-derived osteoblasts/osteocytes as well as patient-derived metastatic cells, further advancing its utility in studying diverse therapeutic responses and guiding individualized treatment strategies.

## 2. Results and Discussion

### 2.1. Experimental Setup and pH Impact of Plasma Treatment

The experimental setup is illustrated in Figure 1A, where the key components, including the XYZ-moving stage, USB spectrometer, plasma source, argon gas supply, and well plate, are schematically depicted. This is complemented by Figure 1B, which captures a photograph of the plasma treatment in action, focusing on the plasma source in detail. To provide a dynamic perspective, an accompanying video (Figure S1) demonstrates the automation of plasma treatment across a well plate, showing the computer-controlled XYZ-moving stage precision of the process. Together, these visuals and dynamic perspectives establish a comprehensive overview of the experimental system and its capabilities.

**Figure 1.**
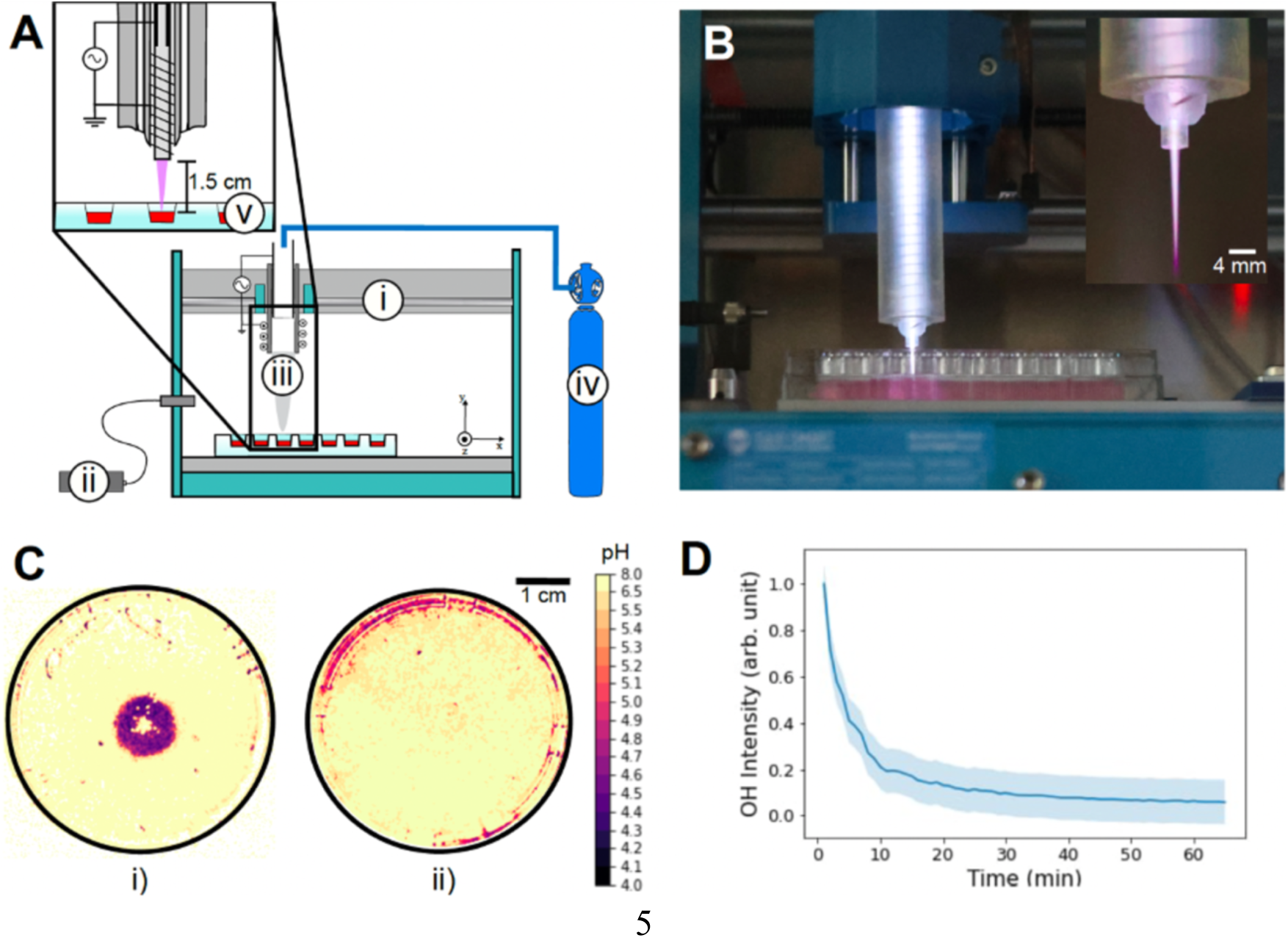
Plasma Treatment Setup. A) Schematic of the experimental setup, including i. an XYZ-moving stage, ii. a USB spectrometer, iii. a plasma source, iv. an argon gas supply, and v. a well plate. B) Photograph of the plasma treatment in action, including a close-up view of the plasma source. C) pH Distribution of plasma-treated A1G7 hydrogel in a 48-well plate: i. direct interaction with the hydrogel, and ii. liquid media layer on top of the A1G7 hydrogel. D) Stabilization time of the OH intensity spectrum during the treatment. Error bars are represented as SD and for *n* = 3.

The interaction between plasma treatment and the surrounding environment is further analyzed in Figure 1C, which compares localized pH changes following stationary plasma treatment on A1G7 hydrogel. When plasma interacts directly with an uncovered hydrogel (Figure 1C i), a distinct and localized drop in pH is observed. In contrast, when the hydrogel is submerged under 2 mm of liquid culture medium during treatment (Figure 1C ii), the localized low-pH region disappears, leaving only a slight overall reduction in pH. This highlights the critical role of the liquid-buffered interface -analogous to biological fluids such as blood, which is also buffered-in distributing reactive species more uniformly and enhancing the homogeneity of plasma treatment.

The dynamic behavior of plasma-produced reactive species is further explored in Figure 1D, which shows the temporal evolution of OH optical emission intensity as measured by an optical spectrometer. The OH emission comes from excited hydroxyl radicals mostly generated in the plasma from the feed gas humidity. The humidity diminishes as the gas flow depletes moisture but stabilizes after approximately one hour. Given that the feed gas humidity strongly influences plasma efficacy in cell inactivation,[37] all plasma treatments are conducted post-stabilization to ensure consistency and accuracy. These results highlight the need to account for feed gas properties in experimental setups.

These observations on the influence of the tissue model environment shown in Figure 1C emphasize the crucial role of the liquid interface in modulating the distribution of RONS and the resulting pH shifts during plasma treatment.[38,39] The liquid interface is fundamental in converting gaseous reactive species into their aqueous counterparts, facilitating the solvation and distribution of RONS.[38] Gaseous OH, for example, dissolves and reacts within the liquid phase, contributing to the formation of aqueous RONS, including H₂O₂ and nitrate (NO₃^-^), through complex chemical pathways.[38] These long-lived species accumulate over time and influence biological interactions.[38] Plasma treatment of liquids also leads to decreased pH due to the formation of acidic species, which can enhance the antibacterial effects of the treated liquid.[38,40–42] Additionally, the liquid’s properties, including thickness and composition, significantly impact RONS penetration and concentration profiles. Thin liquid layers can absorb UV radiation from plasma, leading to further RONS generation and limiting penetration into deeper tissues or hydrogels.[43–45] Structural properties, such as hydrogel density, further constrain diffusion, highlighting the interplay between plasma-generated RONS and their surrounding environment.[39,46]

When a liquid layer covers the hydrogel, it modifies the concentration and distribution of RONS, acting as a mediator that influences their penetration and creates a more complex diffusion profile.[39,47] Unlike plasma-generated RONS, which consist of a mixture of highly reactive and short-lived species capable of permeating the hydrogel, a solution containing only H₂O₂ lacks this complexity and demonstrates limited ability to penetrate the hydrogel matrix, depending on its structural properties.[48] Plasma treatment, however, demonstrates the potential to deliver RONS subcutaneously, penetrating tissues to depths of several millimeters.[39] Biological processes, such as cell signaling and fluid flow, may enhance this delivery, broadening its therapeutic applications.[39] Together, these findings emphasize the importance of the liquid interface in shaping the distribution and effects of plasma-generated reactive species. Building on these insights, the experiments in the subsequent section explore the selective cytotoxicity of CAP treatment on cancer and healthy cells in 2D culture models, providing a foundation for evaluating its therapeutic potential.

### 2.2. Selective Viability Effects of CAP in 2D Culture Models

The selectivity of plasma treatment toward cancer cells over healthy cells is thoroughly demonstrated in Figure 2, which focuses on identifying effective treatment distances and durations while assessing the initial selectivity of CAP. This controlled approach evaluates its selective effects in a simplified system before progressing to more complex 3D culture models. Metabolic activity results as a function of treatment time demonstrate a stronger/faster reduction in cancer cells compared to healthy cells, confirming the selective targeting capability of plasma treatment. As illustrated in Figures 2A, Figure S2A, and Figure S3A, cancer cell metabolic activity decreases significantly with longer treatment durations, while healthy cells maintain relatively higher levels of metabolic activity (% to control). This pattern is evident as early as 1-day post-treatment (Figure 2A) and persists across the 2-(Figure S2A) and 3-day (Figure S3A) observation periods.

**Figure 2.**
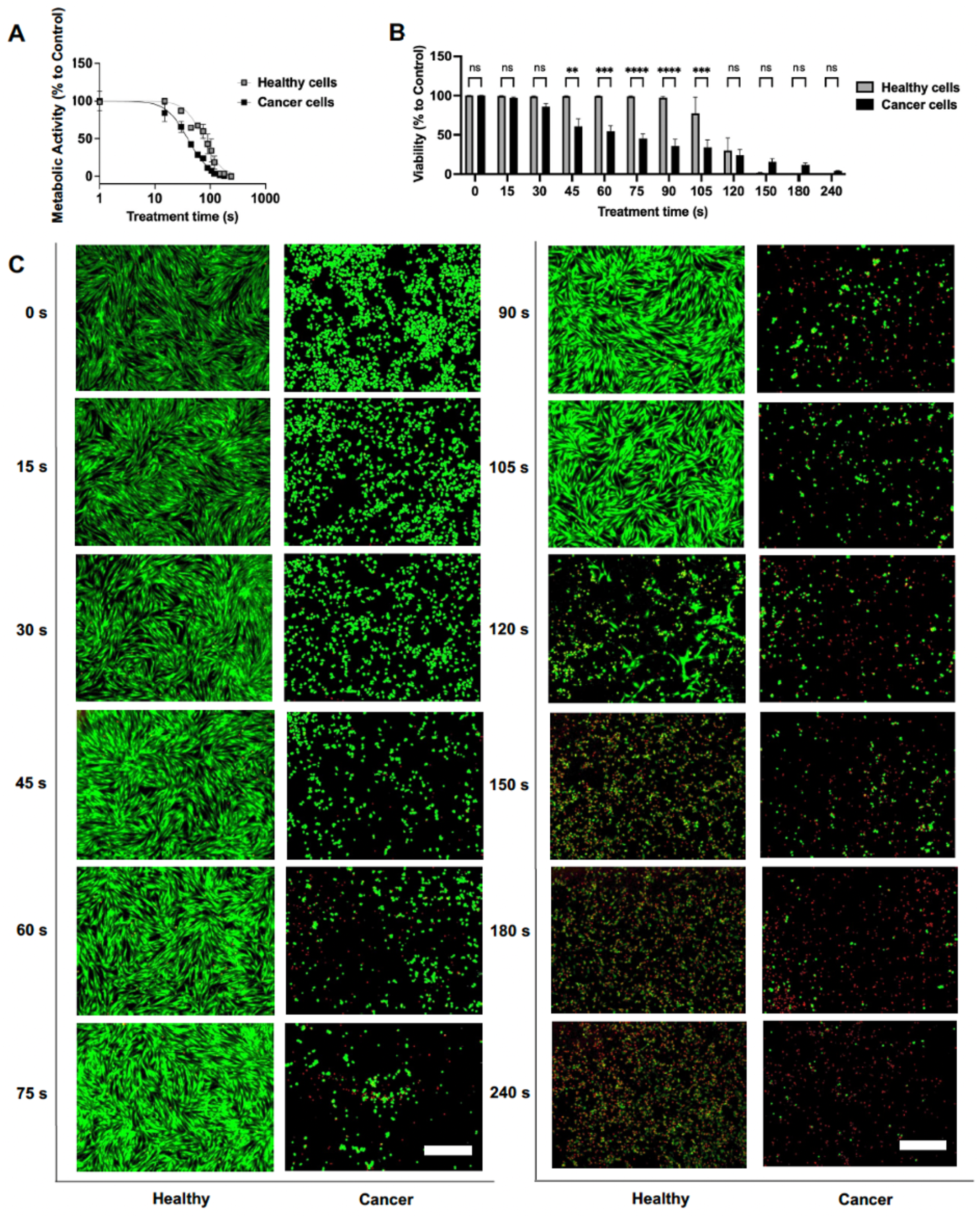
Treatment time-dependent effects of CAP on metabolic activity and viability of 2D cultured cancer and healthy cells 1-day post-treatment. A) Time response of the metabolic activity (% compared to control) of cancer and healthy cells for one day after plasma treatment of different durations. B) Viability (% compared to control) of cancer and healthy cells for 1 day after plasma treatment of different durations. C) Live/Dead microscopic images (Green: Calcein-AM indicating live cells; Red: Ethidium homodimer-1 indicating dead cells) of the viability of the related cancer and healthy cells from (B), with a scale bar of 750 μm. Statistical significance was determined using two-way ANOVA followed by the Šidák multiple comparison test. Error bars are represented as SEM. Statistical significance is indicated as follows: **** *p* < 0.0001, *** *p* < 0.001, ** *p* < 0.01, * *p* < 0.05; *n* = 3.

Cell viability data (quantified from Live/Dead images) provides further confirmation of these selective effects. One day after plasma treatment, 45 to 105-second durations significantly reduced cancer cell viability, as shown in Figure 2B. Healthy cells, in contrast, exhibit only a slight decrease in viability under the same conditions. The greatest selectivity is observed for 90 seconds of plasma exposure 1-day post-treatment, with a 61.45% difference in viability between cancer and healthy cells (p < 0.0001). This trend continues over time, with selectivity towards cancer cell death observed for 60 to 150 seconds of treatment after two days (Figure S2B). By day three, cancer cells still exhibit significant reductions in viability at durations of 90 to 150 seconds, while healthy cells remain largely unaffected (Figure S3B).

Fluorescence microscopy imaging further supports these findings. As shown in Figure 2C, Live/Dead staining images taken one day after treatment visually confirm the quantitative results, highlighting more pronounced damage in cancer cells compared to healthy cells.

These visual differences persist over time, as evidenced by images taken two and three days after treatment (Figure S2C, Figure S3C), providing additional evidence of plasma’s selective cytotoxic effects.

Building on these initial observations, Figure 3 shows the dynamic response of both cell types over time. This analysis explores recovery trends and adaptation signs at 1-, 2-, and 3-day intervals post-treatment, providing insights into the temporal evolution of selective cytotoxicity and potential signs of resistance.

**Figure 3.**
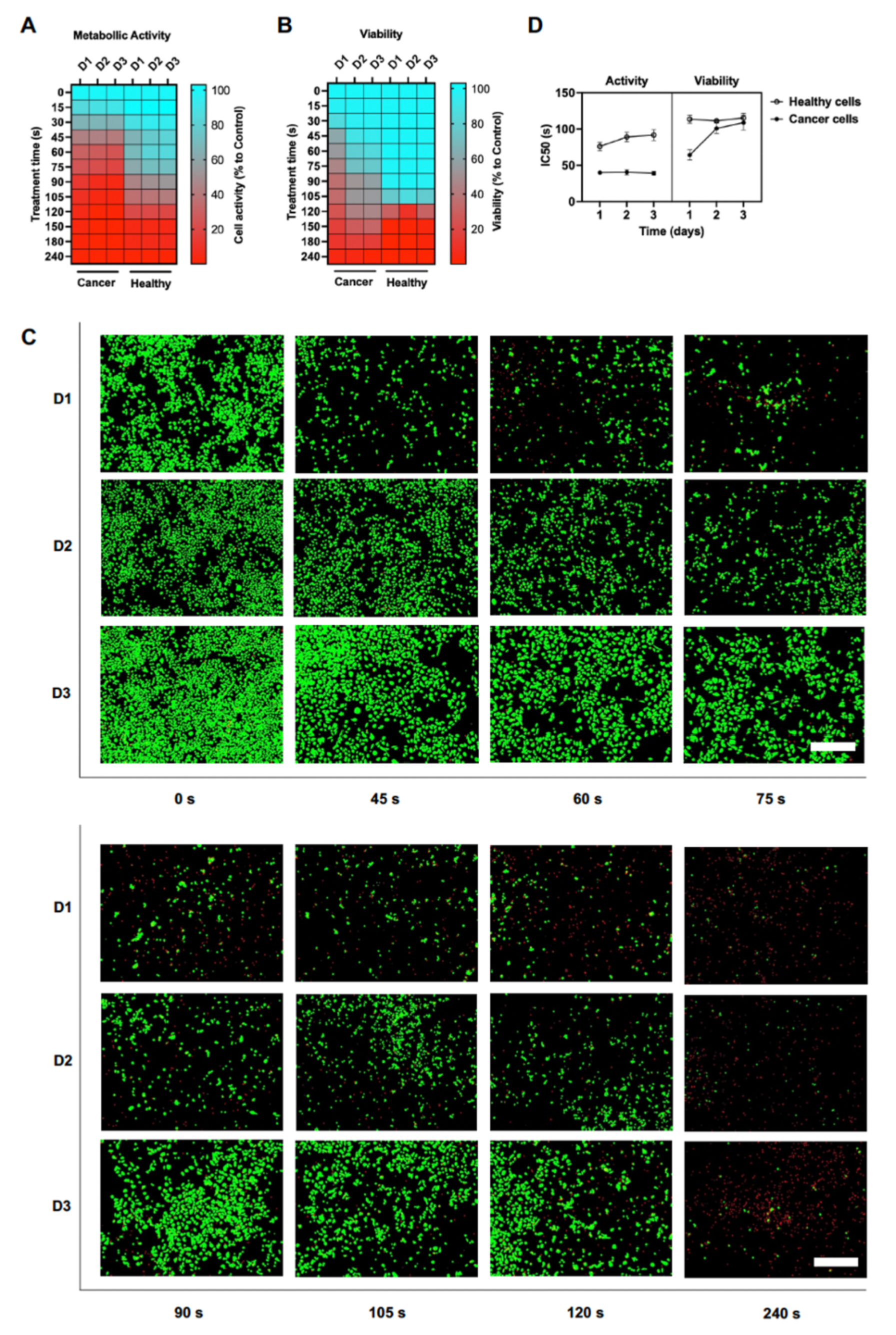
Impact of CAP treatment on metabolic activity and viability of 2D cultured cancer and healthy cells over time. A) Heatmap of the metabolic activity (% compared to control) of cancer and healthy cells for 1, 2, and 3 days after plasma treatment of different durations. B) Heatmap of the viability (% compared to control) of cancer and healthy cells for 1, 2, and 3 days after plasma treatment of different durations. C) Live/Dead microscopic images (Green: Calcein-AM indicating live cells; Red: Ethidium homodimer-1 indicating dead cells) of the viability of the cancer cells for 1, 2, and 3 days after plasma treatment, with a scale bar of 750 µm. D) IC50 of the Time response curves of the metabolic activity and viability (% compared to control) of cancer and healthy cells for 1, 2, and 3 days after plasma treatment of different durations. Error bars are represented as SEM and *n* = 3.

The heatmap of metabolic activity after 1, 2, and 3 days of plasma treatment (Figure 3A) shows that metabolic activity remains relatively stable for both cancer and healthy cells throughout the 3 days, with no significant differences in response to the treatment. This suggests that plasma exposure does not drastically affect the metabolic function of either cell type from 1 to 3 days post-CAP treatment.

In contrast, the Live/Dead assessment of the cell viability heatmap (Figure 3B) reveals a significant increase in viability for cancer cells treated for 45 to 120 seconds from day 1 to day 3 (Figure S4A). This observation indicates that the plasma dose, in the present case, defined by treatment duration and number of treatments, may not have been sufficient to induce complete cell death of all cancer cells within these treatment durations. The remaining cells likely survived or resisted the initial stress and resumed proliferation over time, leading to an overall increase in cell numbers and, hence, detected viability ratio. These findings emphasize the need to optimize plasma treatment parameters to achieve sustained cytotoxic effects, particularly for durations that do not deliver a lethal dose to the entire cancer cell population. A single plasma treatment may only partially affect cancer cells, allowing a subpopulation to survive, proliferate, and possibly take on a therapy-resistant maladaptive phenotype. To address this, introducing repeated plasma treatments at optimized intervals enhances cancer cells’ cytotoxicity and prevents regrowth. The viability of healthy cells remains largely unchanged across the same time points and treatment durations, as shown in Figure S4B, further underscoring the selective nature of plasma treatment. Live/Dead microscopic images of the cancer cells provided in Figure 3C visually corroborate quantified findings from Figure 3B. The images show visible damage to cancer cell viability over increasing CAP treatment durations, with cancer cell viability increasing from 1 day to 2 and 3 days post-CAP treatment.

The IC50 values for time responses related to metabolic activity (Figure 3D) remain stable for cancer cells, with no significant changes over time, while a slight increase is observed for healthy cells. Conversely, the IC50 values for viability via the Live/Dead assay are stable for healthy cells but show a slight increase for cancer cells. These findings align with the Live/Dead cell viability heatmap (Figure 3B), reinforcing the conclusion that the plasma dose may not have been sufficient to completely eliminate cancer cells, leaving some cells to recover and proliferate.

These results highlight the complex dynamics of plasma-induced cytotoxicity and selectivity, showing that specific treatment durations may partially affect cancer cells, leaving room for recovery and regrowth while largely sparing healthy cells (Figure S4B). This aligns with findings from other studies demonstrating the efficacy of CAP treatment against MDA-MB-231 breast cancer cells in 2D cultures. For example, CAP treatment has demonstrated greater vulnerability of triple-negative breast cancer cells, such as MDA-MB-231, compared to non-cancerous breast epithelial control cells (MCF10A).[49,50] This selectivity was attributed to the differential accumulation of intracellular ROS, hyperactivated signaling pathways such as Mitogen-Activated Protein Kinase (MAPK)/c-Jun N-terminal Kinase (JNK) and Nuclear Factor kappa-light-chain-enhancer of activated B cells (NF-κB), as well as the inhibition of the interleukin-6 (IL-6) pathway in cancer cells.[49,50] While CAP has also been more effective against MDA-MB-231 cells than against hbmMSCs, the selective cytotoxicity of the treatment heavily relies on its conditions and exposure time.[51] Cancer cell proliferation has been observed to vary depending on CAP treatment duration, with shorter treatments allowing some recovery by day three. However, longer treatment durations, such as 60 and 90 seconds, more effectively suppress proliferation by day three, with stabilization observed by day five.[51]

While most studies have focused on 2D culture models, they provide valuable insights into the mechanisms of plasma-induced effects. These include apoptosis induction, epigenetic changes, and cellular responses such as altered proliferation rates, oxidative stress responses, and changes in signaling pathways. For instance, argon plasma treatments have been shown to induce epigenetic alterations in 2D cultures.[52] Additionally, comparisons of helium and helium-oxygen plasma treatments with standard drug therapies[53,54] and differences in apoptotic protein expression between direct plasma and plasma-activated media (PAM) treatments further highlight the diverse effects of plasma treatment.[54] However, these findings represent a simplified approximation of the cellular environment, as 2D models lack the complexity of *in vivo* conditions. To better capture these complexities, more advanced models, such as 3D culture models, are required. These systems can provide a more comprehensive understanding of plasma effects, including cell-cell interactions, extracellular matrix influences, and the spatial distribution of reactive species. To address this gap, our subsequent experiments aim to explore the effects of plasma treatment in more physiologically relevant 3D culture models.

### 2.3. Selective Viability Effects of CAP in 3D Monoculture Models

The time-response results in 3D monoculture models, where each cell type is bioprinted independently, were analyzed to evaluate the selectivity of plasma treatment for cancer cells over healthy cells and to observe its efficacy over time. These experiments are crucial for validating findings from 2D models in a more physiologically relevant 3D context, allowing for an assessment of whether direct plasma treatment maintains its selective effects under these conditions. Measurements of metabolic activity in 3D monoculture, illustrated in Figure 4A and its supplementary counterparts (Figure S5A and Figure S6A), demonstrate clear selectivity for cancer cells over healthy cells across 1, 2, and 3 days post CAP treatments.

**Figure 4.**
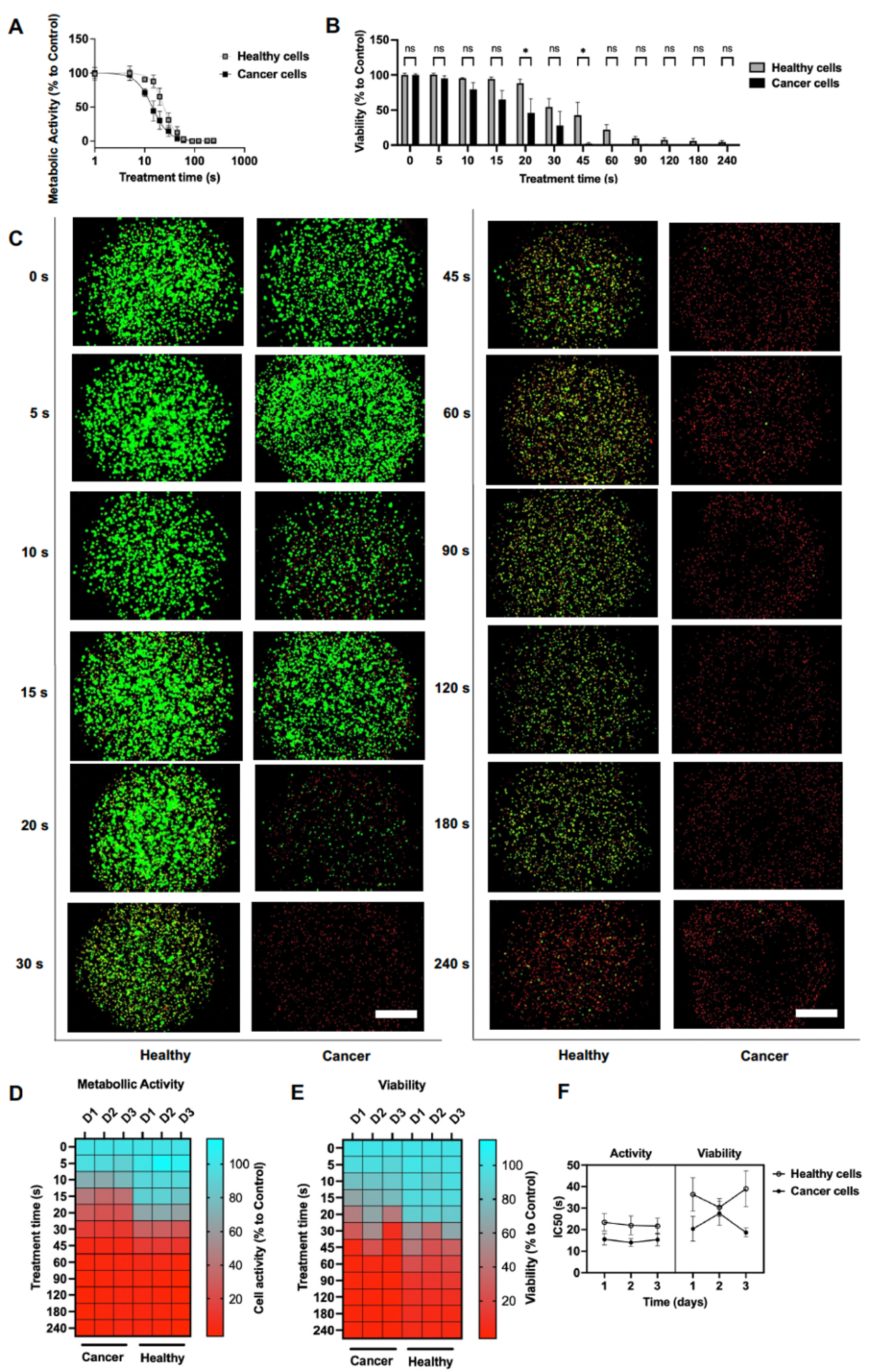
Time-dependent effects of CAP treatment on metabolic activity and viability of 3D monocultures of cancer and healthy cells over time. A) Time response of the metabolic activity (% compared to control) of cancer and healthy cells for 1 day after plasma treatment of different durations. B) Viability (% compared to control) of cancer and healthy cells for 1 day after plasma treatment of different durations. C) Live/Dead microscopic images (Green: Calcein-AM indicating live cells; Red: Ethidium homodimer-1 indicating dead cells) of the viability of the related cancer and healthy cells from (B), with a scale bar of 750 μm. D) Heatmap of the metabolic activity (% compared to control) of cancer and healthy cells for 1, 2, and 3 days after plasma treatment of different durations. E) Heatmap of the viability (% compared to control) of cancer and healthy cells for 1, 2, and 3 days after plasma treatment of different durations. F) IC50 of the time response curves of the metabolic activity and viability (% compared to control) of cancer and healthy cells for 1, 2, and 3 days after plasma treatment of different durations. Statistical significance was determined using two-way ANOVA followed by the Šidák multiple comparison test. Error bars are represented as SEM. Statistical significance is indicated as follows: **** *p* < 0.0001, *** *p* < 0.001, ** *p* < 0.01, * *p* < 0.05; *n* = 3.

Similarly, cell viability data (Figure 4B) reveal significant selectivity (p < 0.05) for 20 and 45 seconds of treatment after 1 day. The greatest selectivity is observed for 20 seconds, with a 42.4% difference in viability between cancer and healthy cells (p < 0.05). However, by day 2, statistical significance diminishes (Figure S5B), though selectivity re-emerges on day 3 with treatment durations of 20 and 30 seconds (Figure S6B), suggesting dynamic changes in cellular responses over time. Visual confirmation of these trends is provided by Live/Dead microscopic images (Figure 4C), which reveal pronounced differences in viability between cancer and healthy cells 1-day post-treatment. Additional images from days 2 and 3 (Figure S5C and Figure S6C) further highlight the sustained impact of plasma treatment on cancer cells compared to healthy cells, where proliferation is also stunted.

Heatmaps of metabolic activity and viability shown in Figure 4D and Figure 4E, respectively, further support these findings. The heatmaps indicate minimal changes in both cancer and healthy cells over 1, 2, and 3 days, suggesting that plasma exposure does not drastically alter metabolic function or overall viability. These non-significant viability changes are quantified in Figure S7 for both cell types at 1-, 2-, and 3-day post-treatment. The stability observed in the 3D models contrasts with the regrowth seen in 2D cultures, emphasizing the value of 3D systems in better mimicking physiological conditions. This difference can be partly attributed to the structural complexity of the 3D matrix. Studies have shown that higher plasma treatment durations are required to effectively penetrate and treat 3D structures than 2D cultures, where cells are exposed more directly to plasma.[55] Additionally, the lower cell density within the 3D matrix may inherently limit proliferation and regrowth, contributing to the observed stability over time. IC50 values for metabolic activity and viability (Figure 4F) consistently demonstrate lower thresholds for cancer cells than healthy cells across all treatment conditions, underscoring plasma treatment’s selectivity. Moreover, the IC50 values remain stable from day 1 to day 3 within the error margins, highlighting the consistency of these effects over time.

While several studies[56,57] have explored PAM in 3D models, they differ significantly from direct plasma treatments. For instance, PAM studies have shown synergistic effects when combined with standard drug therapies in spheroid models and demonstrated selective action against triple-negative breast cancer cells.[56] However, the mechanisms and dynamics of direct plasma treatment in 3D environments for triple-negative breast cancer cells remain largely underexplored, pointing to a critical gap in the literature. Our findings help bridge this gap by providing new evidence, demonstrating for the first time, to our knowledge, the selectivity of direct plasma treatment for triple-negative breast cancer cells over healthy cells in bioprinted tissue-like constructs. These results represent a significant step in leveraging bioprinted constructs to investigate selective plasma action mechanisms and optimize treatment parameters in more complex 3D environments.

Building on these findings, we provide, to our knowledge, the first 3D co-culture model that incorporates both cancer and healthy stromal cells within a bioprinted construct for investigating CAP treatment on cancer cells. This innovative approach enables the investigation of plasma treatment selectivity and effectiveness in a more physiologically relevant system that closely mimics the tumor microenvironment, including cell-cell interactions and tissue heterogeneity. These advancements pave the way for refining plasma therapies and improving their translational potential for clinical applications.

### 2.4. Selective Viability Effects of CAP in 3D Co-Culture Models

The selective effects of CAP treatment in a 3D bioprinted co-culture model designed to mimic a complex tumor environment are evaluated in Figure 5. This analysis compares CAP treatment selectivity in targeting cancer cells over healthy cells with the standard chemotherapeutic agent doxorubicin, highlighting CAP potential as a localized treatment in a physiologically relevant 3D context. The viability results clearly demonstrate that CAP treatment selectively targets cancer cells over healthy cells in a 3D bioprinted co-culture model, where cancer cells are positioned in the center and are surrounded by healthy cells.

**Figure 5.**
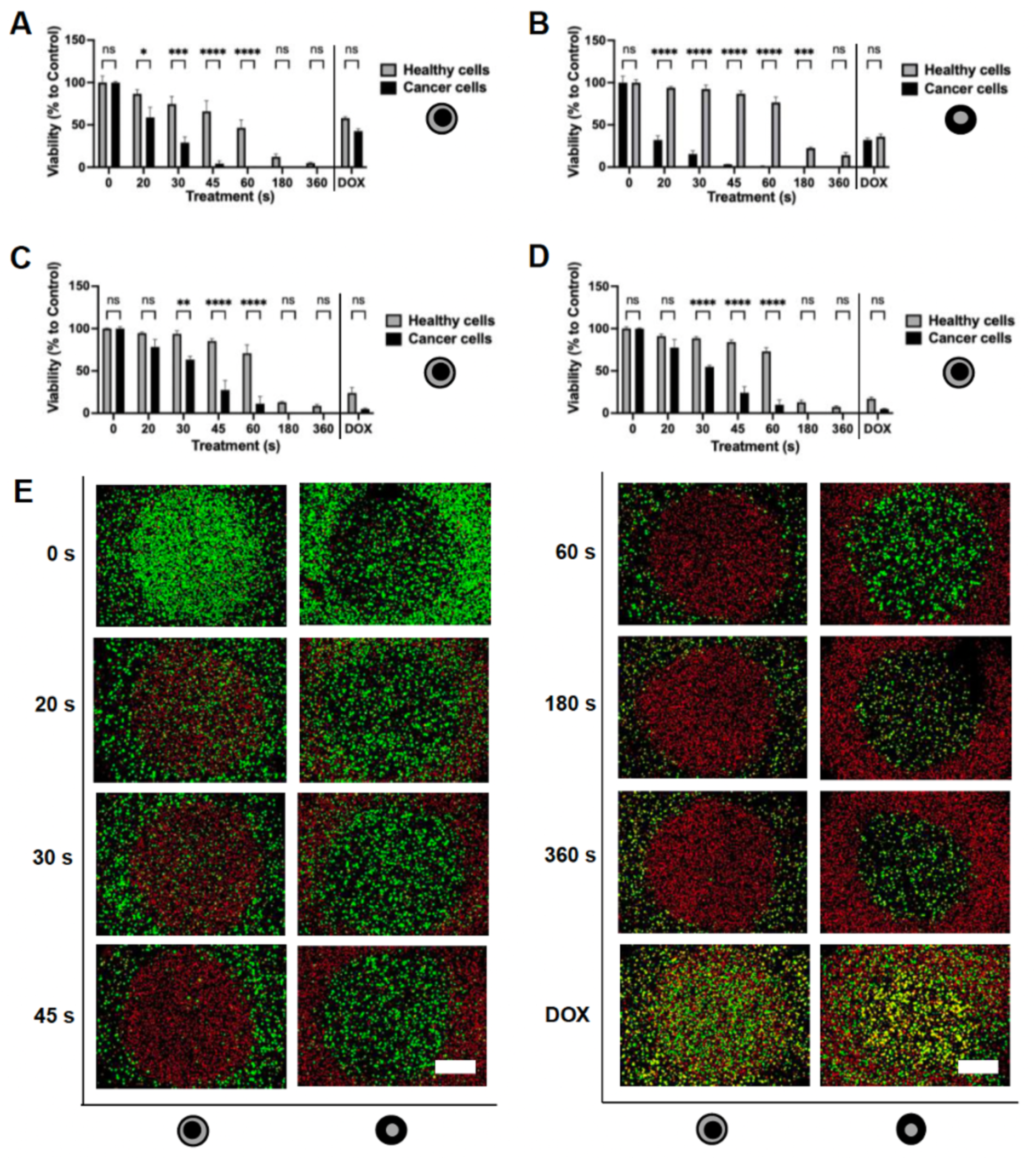
Viability evaluation of 3D bioprinted co-culture model after CAP treatment over time through Live/Dead assay. A) Viability (% compared to control) of the co-culture model (cancer cells positioned in the center and healthy cells surrounding them) one day after CAP treatment of different durations, compared to doxorubicin treatment (5 µм). B) Viability (% compared to control) of the reversed co-culture model (healthy cells positioned in the center and cancer cells surrounding them) one day after CAP treatment of different durations, compared to doxorubicin treatment (5 µм). C) Viability (% compared to control) of the co-culture model 2 days after CAP treatment of different durations, compared to doxorubicin treatment (5 µм). D) Viability (% compared to control) of the co-culture model three days after CAP treatment of different durations, compared to doxorubicin treatment (5 µм). E) Live/Dead microscopic images (Green: Calcein-AM indicating live cells; Red: Ethidium homodimer-1 indicating dead cells) of cell viability in the co-culture model and the reversed co-culture model one day after CAP treatment of different durations, compared to doxorubicin treatment (5 µм). Scale bar = 1250 µm. Statistical significance was determined using two-way ANOVA followed by the Šidák multiple comparison test. Error bars are represented as SEM. Statistical significance is indicated as follows: **** *p* < 0.0001, *** *p* < 0.001, ** *p* < 0.01, * *p* < 0.05; *n* = 3.

Significant selectivity is observed for 20 and 60 seconds of treatment on day 1, as cancer cell viability decreases significantly more than that of healthy cells, whereas doxorubicin treatment indiscriminately kills both cancer and healthy cells (Figure 5A). The greatest plasma selectivity is observed for 45 seconds of treatment, with a 61.70% difference in viability between cancer and healthy cells (p < 0.0001). To confirm that geometric factors did not influence these results, the configuration was reversed, with healthy cells at the center and cancer cells surrounding them. The results confirmed the selectivity of CAP treatment against cancer cells, with significant effects observed after 20 and 180 seconds of treatment, while doxorubicin again showed indiscriminate effects (Figure 5B). The greatest selectivity in the reversed model is observed for 45 seconds of treatment, with an 83.60% difference in viability between cancer and healthy cells (p < 0.0001).

The selectivity of CAP treatment persisted over subsequent days. In the standard co-culture model, with cancer cells in the center, significant selectivity was observed on day 2 for 30 to 60 seconds of treatment, with the greatest selectivity for 60 seconds resulting in a 59.63% difference in viability between cancer and healthy cells (p < 0.0001, Figure 5C). On day 3, similar results were observed, with the greatest selectivity again for 60 seconds, showing a 62.98% difference in viability (p < 0.0001, Figure 5D). Throughout the 3 days, doxorubicin continued to show non-selective effects in either configuration. Visual evidence from Live/Dead fluorescence microscopic images (Figure 5E) corroborates these findings, revealing more pronounced effects of CAP treatment on cancer cells than on healthy cells.

Further analysis of viability over 1, 2, and 3 days highlights that CAP significantly reduces cancer cell viability with 20, 30, and 45 seconds of treatment time, while healthy cell viability is only marginally affected at higher durations, such as 30, 45, and 60 seconds. In contrast, doxorubicin’s effects on viability remain non-selective across all conditions (Figure S8A–C).

The observed selective effects of CAP treatment can be attributed to the generation of RONS, such as hydrogen peroxide (H₂O₂), hydroxyl radicals (OH), and nitrogen species like peroxynitrite (ONOO^-^), which play key roles in inducing selective cytotoxicity in cancer cells.[58,59] These RONS disrupt mitochondrial functions by impairing the electron transport chain, leading to oxidative stress, loss of mitochondrial membrane potential, and the release of pro-apoptotic factors such as cytochrome c, which activate caspase-dependent apoptotic pathways.[60] RONS also promote lipid peroxidation by attacking unsaturated fatty acids in cellular membranes, generating lipid radicals and reactive aldehydes like 4-hydroxynonenal (4-HNE). This oxidative damage compromises membrane integrity and can induce ferroptosis, a regulated cell death pathway dependent on iron and lipid peroxidation.[61] Furthermore, direct CAP treatment may induce pyroptosis - a pro-inflammatory form of cell death - through pathways that activate caspase-1 and the subsequent cleavage of gasdermin D. Both direct and indirect CAP treatment also trigger apoptosis and ferroptosis through the generation of reactive species and subsequent oxidative damage.[62] These mechanisms highlight CAP treatment versatility in targeting cancer cells, which are particularly vulnerable due to their altered metabolism and elevated reactive species levels.[62,63] In contrast, healthy cells, such as hbmMSCs, exhibit more robust antioxidant defenses, including higher levels of glutathione and catalase, which mitigate oxidative damage and protect them from RONS-induced cytotoxicity.[64,65]

Unlike CAP, doxorubicin - a widely used chemotherapeutic for breast cancer - primarily works by intercalating DNA and inhibiting topoisomerase II, leading to the generation of free radicals, subsequent DNA damage, and apoptosis.[66] While doxorubicin’s mechanism involves the production of reactive species, its impact is less targeting than plasma-generated RONS.[67] This lack of specificity impacts both cancerous and healthy cells, causing systemic toxicity, including cardiotoxicity and other side effects.[66] Consequently, dose reductions are often necessary, potentially compromising treatment efficacy.[66] CAP treatment, by contrast, offers a more localized therapeutic approach. By generating RONS with spatial precision, CAP enables selective targeting of cancer cells, particularly in 3D cultures, where the spatially confined effects of plasma minimize damage to surrounding healthy tissues.[58,59] The ability of CAP to selectively target cancer cells arises from the differing vulnerabilities of cancer cells and healthy cells to oxidative stress.[68]

The therapeutic potential of CAP is particularly evident in post-surgical applications. After tumor removal, CAP can be applied directly to the surgical site to target residual cancer cells, reducing the risk of recurrence while minimizing systemic toxicity. CAP could be integrated into clinical workflows as an intraoperative or minimally invasive therapy for treating bone metastases, using portable plasma devices to target challenging anatomical regions like the spine or pelvis.[10] However, scaling CAP for broader clinical use requires the development of standardized, cost-effective devices and optimized treatment protocols to ensure reproducibility across different clinical settings, as well as to account for variation in tissue composition and anatomical complexity.[69] Integrating CAP into existing surgical tools, such as robotic or endoscopic systems, could further enhance its accessibility and practicality.[70,71] While CAP may not entirely replace conventional drug therapies like doxorubicin, its localized action and ability to minimize systemic toxicity make it a promising strategy for specific clinical applications, particularly in post-surgical and targeted treatments.

Advanced 3D co-culture models are essential for studying CAP treatment mechanisms in physiologically relevant settings. Commonly studied 3D co-culture models include spheroids,[55,72,73] organoids,[74] and scaffolds,[58] which are widely used to replicate the tumor microenvironment *in vitro*. While these models provide valuable insights into tumor biology and treatment responses, they often lack the complexity needed to accurately mimic the cellular interactions, spatial arrangements, and vascular dynamics present in real tumors.[75] These limitations highlight the potential of more advanced systems, such as bioprinted models, which provide opportunities to study complex cell interactions in a controlled environment. Bioprinted models offer significant advantages by allowing customizable architectures that mimic tissue organization and incorporating features like stromal components and multicellular arrangements.[29,33,34] Their physiological relevance is supported by features such as hypoxia gradients, nutrient diffusion, and structural integrity, which reflect key aspects of the tumor microenvironment.[76] Future innovations, such as integrating dynamic perfusion systems and immune components, could further enhance their ability to mimic *in vivo* conditions, broadening their applicability to CAP.[33,34,77]

Despite these advancements, no prior studies have employed bioprinted co-culture 3D models to investigate the effects of CAP treatment on MDA-MB-231 cancer cells. To the best of our knowledge, this study is the first to introduce a bioprinted co-culture model specifically designed to study the impact of plasma treatment on cancer cells, offering a more sophisticated and physiologically relevant approach to understanding CAP treatment selective effects. While our bioprinted models bridge a critical gap between traditional 2D cultures and animal studies, we acknowledge certain limitations. For instance, it does not incorporate immune cells, assess inflammatory responses, or include blood vessels or fluid flow conditions. While the present study did not evaluate the effects of CAP treatment on cell signaling or apoptosis pathways, our work represents a foundational step in developing a reliable platform that links experimental redox-based therapies, such as CAP treatment, to physiologically relevant models. Our model establishes a proof-of-concept that can be adapted for future studies to explore these mechanisms more deeply. By refining and validating this platform, it will become a valuable tool for advancing CAP research and studying complex therapeutic mechanisms.

## 3. Conclusion

In conclusion, this study is the first to explore the effects of CAP therapy on a 3D bioprinted tumor-stroma model, demonstrating significant selectivity for cancer cells in both 2D and 3D cultures. The selectivity is maintained over multiple days of treatment, although there is a noted increase in cancer cell viability after extended periods. Unlike systemic treatments such as doxorubicin, which targets highly proliferative cells, CAP offers a more localized therapeutic approach, which has the potential to reduce side effects typically associated with systemic therapies. The 3D bioprinted co-culture model provides a platform to study plasma-tissue interactions and tumor-stroma dynamics and investigate the biological pathways through which CAP acts on tissues. This model holds substantial therapeutic promise and can be adapted for future research exploring various cancer types, treatment durations, and regimes. Additionally, incorporating different stromal cell types, immune components, and bioprinted vasculature with endothelial cells offers a versatile tool to investigate further cancer cell resistance, adaptation mechanisms, and the immunological aspects of therapy.

These findings contribute to advancing the understanding of plasma-tissue interactions and developing more effective, targeted, and selective cancer therapies.

## 4. Experimental Section

### Cells and Materials

Human bone marrow-derived mesenchymal stromal cells (hbmMSCs) were obtained from RoosterBio, USA. Triple-negative breast cancer cells (MDA-MB-231) were donated by Dr. Morag Park’s laboratory at McGill University. Phosphate-buffered saline (PBS, Gibco, #10010049), high glucose Dulbecco’s Modified Eagle Medium (DMEM, 4.5 g L^-1^ D-Glucose, Gibco, #12430062), and low glucose DMEM (1 g L^-1^ D-Glucose, #11054020), Penicillin-Streptomycin (PS) (Gibco, #15140122), fetal bovine serum (FBS) (Gibco, #12483020), 0.25% Trypsin-EDTA (Gibco, #25200072), Live/Dead Viability/Cytotoxicity Kit (Invitrogen, #L3224), and AlamarBlue Cell Viability Reagent (Invitrogen, #DAL1100) were purchased from ThermoFisher Scientific, USA. Recombinant Human Fibroblast Growth Factor 2 (FGF2, 154 a.a., #100-18B-100UG) was obtained from PeproTech, USA. T75 (#83.3911.002) and T175 (#83.3912.002) flasks, as well as 48-well standard plates (#83.3923), were from Starstedt, Germany. The 48-well suspension culture plates (Cellstar, Greiner Bio-One, #82051-000) for 3D cultures were obtained from VWR, USA. 96-well plates (Corning, half-area microplate, flat bottom clear, #CLS3882), calcium chloride dihydrate (#223506), doxorubicin hydrochloride (#44583), and gelatin powder from bovine skin (#69391-500G) were purchased from Sigma-Aldrich, USA. Sodium alginate powder (Protanal® LF 10/60 LS) was purchased from FMC BioPolymer, USA.

### Hydrogel preparation

The hydrogel, composed of A1G7 with 1 wt% sodium alginate and 7 wt% gelatin, was prepared using low glucose DMEM (1 g L^-1^). The prepared solution was exposed to UV light overnight for sterilization. DMEM was chosen over regular PBS or DI water for making the hydrogel due to its role in supporting cell viability and function during 3D bioprinting.[78] Additionally, low glucose DMEM with 1 g L^-1^ was preferred over high glucose DMEM (4.5 g L^-1^) because it more closely mimics human physiological conditions. The human body typically regulates blood glucose levels to range from 0.72 to 1.08 g L^-1^ to maintain homeostasis.[79] The hydrogel solution was then stored in a refrigerator at 4 °C and used within 1 month.

### Cell culture preparation

MDA-MB-231 and hbmMSCs cells were cultured in high glucose DMEM supplemented with 10% FBS and 1% PS. For hbmMSCs, FGF2 was added at a concentration of 2 µL per 40 mL (100 µg mL^-1^ aliquots). MDA-MB-231 cells were grown in T75 flasks, while hbmMSCs were grown in T175 flasks at 37 °C and 5% CO2. Once 80% confluence was reached, cells were washed with PBS and harvested using 0.25% trypsin-EDTA. Only low passage hbmMSCs (passage number p2) were used for experiments, and MDA-MB-231 cells with passage numbers 13 to 23 were employed. For 2D cultures, MDA-MB-231 and hbmMSCs cells were seeded in 48-well standard plates at a density of 35,000 cells per well, using high glucose DMEM supplemented with 3% FBS and 1% PS. The plates were incubated at 37 °C and 5% CO2 for 24 hours before automated CAP treatment in a total media volume of 500 µL. For 3D bioprinting, MDA-MB-231 and hbmMSCs cells were mixed into a sterile hydrogel at a density of 2 × 10^6^ cells mL^-1^. The hydrogel was preheated in a water bath at 37 °C to achieve thermal equilibrium and a liquid-like consistency. Air bubbles introduced during the mixing process were removed by centrifugation for 1 minute at 1500 rpm (Benchtop Centrifuge, #Q5120, ABM). Cell-laden hydrogel syringes were then incubated (37 °C, 5% CO2) for 30 minutes before bioprinting.

### Bioprinting Design and Parameters

A circulator co-culture model was designed using Tinkercad (Autodesk, USA), as shown in Figure S9. It features an inner disk with a diameter of 2.6 mm and a height of 1 mm, which contains MDA-MB-231 cancer cells embedded in A1G7 hydrogel, and an outer ring with a diameter of 4.4 mm and a height of 1 mm, containing hbmMSCs healthy cells also embedded in A1G7 hydrogel. The CAD (computer-aided design) model was converted to STL (stereolithography) format and sliced with PrusaSlicer version 2.8.1 (Prusa Research, Czech Republic) to generate a G-code. This G-code was manually modified to print the outer disk layers first and the inner ones last to prevent the automated centering of the cartridge from crushing the construct. Bioprinting was performed with the Bio X 3D bioprinter (Cellink, Sweden), equipped with up to three printheads. The cell-laden hydrogel was incubated for 30 minutes (37 °C, 5% CO2) and then placed at room temperature in the hood for 15 minutes to achieve optimal bioprinting rheology. For monocultures, the 3D bioprinted constructs (inner disk diameter 2.6 mm, height 1 mm) were printed using a temperature-controlled printhead with sterile conical 22G bioprinting nozzles from Cellink. Printing parameters included a temperature of 26 °C, pressure of 3 to 9 kPa for MDA-MB-231 and 4 to 18 kPa for hbmMSCs cells, and a speed of 8 mm s^-1^. For co-cultures, two pneumatic BioX printheads with conical 22G nozzles were used in 48-well suspension culture plates, with printing parameters of 5 to 10 kPa for MDA-MB-231 and 16 to 25 kPa for hbmMSCs cells, and a speed of 8 mm s^-1^. After bioprinting, the constructs were chemically crosslinked by immersing them in a 0.1 м calcium chloride (CaCl2) solution for 5 minutes. The CaCl2 solution was prepared by dissolving CaCl2 dihydrate in MilliQ water. The constructs were then washed with PBS and incubated with 500 µL of high glucose DMEM supplemented with 1% FBS and 1% PS at 37 °C and 5% CO2 for 24 hours before automated CAP treatment (for both monocultures and co-cultures) and doxorubicin control treatment (for co-cultures). The doxorubicin drug was mixed with 500 µL of low serum media (high glucose DMEM supplemented with 1% FBS and 1% PS) to achieve a concentration of 5 µм.

### Plasma source

The plasma source used in this experiment is a custom-built cold atmospheric pressure plasma jet inspired by a coaxial dielectric barrier discharge (DBD) design (see Figure 1). The device consists of a stainless-steel cylinder (OD = 0.078”, ID = 0.063”, McMaster-Carr) as the high-voltage electrode housed within a quartz tube (OD = 4 mm, ID = 2 mm, Thomas Scientific). The high-voltage electrode is positioned 10 cm away from the jet nozzle, with a copper coil (gauge 24) wrapped around the quartz tube as the ground electrode, spaced 5 mm between loops. The power supply (PVM500-1000L, Information Unlimited, USA) delivers 3 kVpp at a resonant frequency of 22 kHz, with a 40% duty cycle applied to regulate plasma generation. High-purity argon gas (99.998%, MEGS) flows through the jet at 2 SLM, surrounded by a gas curtain to stabilize the plasma effluent. The electrical structure is housed in a high-temperature resistant resin (High Temp, FormLabs). The shell was fabricated using stereolithography 3D printing, comprising two detachable sections (support and cap) for electrical maintenance access. The jet’s design integrates directly with a computer-controlled XYZ-moving stage system (Genmitsu 3018-PROVer, SainSmart, USA), enabling automated positioning and plasma ignition. The XYZ-moving stage was modified to use a TTL signal to trigger the power supply for plasma activation. For biological tests, the XYZ-moving stage was programmed to position the jet in each well at a fixed plasma position, ensuring precise control of the 1.5 cm gap between the plasma source and the media surface and accurate treatment durations for 2D and 3D cultures (see Figure 1).

### pH Measurements

The pH measurements were conducted using sodium fluorescein (Sigma-Aldrich, USA), calibrated with a pH probe (LabSen 241-6 Glass-body pH Electrode, Apera) and an acquisition system (Mi 150, Martini Instruments). A calibration curve was created using a hydrogel solution (1% alginate, 7% gelatin) mixed at six different concentrations with HCl (Sigma-Aldrich, USA). Experimental parameters for direct plasma treatment included a distance of 1.5 cm and a gas flow rate of 2 SLM of argon. The methodology was based on the work of Busco et al.[80] A calibration curve was developed to link fluorescence intensity to pH by correlating the degradation of fluorescein with pH decrease. Due to its pH sensitivity and established use in angiography, fluorescein was selected as a reliable choice for tissue-related applications.[81] For imaging fluorescence, a 25 mW, 405 nm Violet Alignment Laser was used as the excitation source. The emitted fluorescence was captured using a CMOS camera (Allied Vision Alvium 1800 U-240m, Edmund Optics, USA) and a 475 nm long pass optical filter (Edmund Optics, USA).

### Metabolic Activity Assay

The metabolic activity of MDA-MB-231 and hbmMSCs cells was measured using the AlamarBlue Cell Viability Reagent kit on days 1, 2, and 3, following the manufacturer’s instructions. For each analysis day, media and treatments were removed and stored in the incubator (37 °C, 5% CO2). A solution of 10 % AlamarBlue and 90 % low-serum fresh media was prepared, with 3% FBS and 1% PS for 2D cultures and 1% FBS and PS for 3D cultures. The solution was then added to the wells, with incubation at 37 °C and 5% CO2 for 4 hours for 2D cultures and 6 hours for 3D cultures. After incubation, 100 µL of the solution was pipetted into 96-well plates and analyzed using a Tecan Infinite M200 Pro microplate reader (Tecan, Switzerland) at an excitation wavelength of 560 nm and an emission wavelength of 590 nm to record metabolic activity. Once the AlamarBlue solution was removed, the wells were washed with PBS, and the original control and treated media were reapplied to continue treatment for Days 2 and 3. The metabolic activity was then reassessed on these days. The experiments were performed in triplicates with *n* = 3 for each cell type on Days 1, 2, and 3. Day 1 corresponds to 24 hours of CAP treatment, Day 2 to 44 hours of treatment in 2D culture and 42 hours in 3D monoculture, and Day 3 to 64 hours of treatment in 2D culture and 60 hours in 3D monoculture. This timing aligns with the incubation requirements of AlamarBlue. Metabolic activity values were adjusted by subtracting the average background signal from the AlamarBlue solution mixed with media without cells. For each condition, the values were normalized to the control by dividing the measured value by the average value of the control and multiplying the result by 100 to express it as a percentage. The normalized values were then averaged across triplicates, and statistical analysis was performed for *n* = 3.

### Cell Viability Assay

The viability of MDA-MB-231 and hbmMSCs cells was assessed using the Live/Dead Viability/Cytotoxicity Kit on days 1, 2, and 3, following the manufacturer’s instructions. Briefly, media was removed from the wells and replaced with 200 µL of Live/Dead solution diluted in PBS. The plates were then incubated at 37 °C and 5% CO2 for 15 minutes for 2D cultures and 30 minutes for 3D cultures before imaging with a digital inverted fluorescence microscope (EVOS M5000, Invitrogen, #AMF5000). Day 1 corresponds to 24 hours of CAP treatment. Day 2 corresponds to 44 hours of treatment in 2D cultures and 42 hours in 3D monocultures. Day 3 corresponds to 64 hours of treatment in 2D cultures and 60 hours in 3D monocultures. This timing aligns with the metabolic activity assessment using AlamarBlue, which required 4 hours of incubation for 2D and 6 hours for 3D cultures. For 3D co-cultures, Day 1 corresponds to 24 hours of treatment, Day 2 to 48 hours, and Day 3 to 72 hours, as only viability was assessed. The images from 2D and 3D cultures were then analyzed as follows: 1 square of area 1.21 mm² per image for 2D cultures, three squares of area of approximately 0.85 µm² per image for 3D monocultures and the inner part of 3D co-cultures, and 1 square of area of approximately 0.85 mm² per image for the outer part of 3D co-cultures (see Figure S10). Cells were counted in the red channel for dead cells and in the merged image for both red and green channels. Cells with overlapping green and red signals were classified as live or dead based on the intensity of the merged channels, with stronger red intensity indicating dead cells and stronger green intensity indicating live cells. Squares for analysis were created using a custom Python code available upon request.

Cell counts were performed using ImageJ2 version 2.1.0 (Fiji, USA). To assess cell viability, the number of live cells was divided by the total cell count and multiplied by 100. For each experimental condition, the viability was normalized to the control by dividing the viability percentage of the condition by the average viability percentage of the control and multiplied by 100. The normalized values were then averaged across triplicates, and statistical analysis was performed for *n* = 3. All microscopic images presented in this manuscript were uniformly enhanced in intensity using the same parameters with ImageJ2 software.

### Statistical Analyses

Relative IC50 values were determined using GraphPad Prism by performing nonlinear regression and fitting a sigmoidal curve to normalized data. The IC50, representing the concentration required for 50% inhibition relative to the normalized maximum response, was reported with its 95% confidence interval. Data are presented as mean ± standard deviation (SD) in Figure 1 and as mean ± standard error of the mean (SEM) in all other Figures. Biological experiments were performed in triplicates for *n* = 3. Statistical significance was evaluated using two-way ANOVA with Šidák multiple comparison test. Analyses were conducted using GraphPad Prism version 10.3 (GraphPad Software, USA), with significance levels indicated as \*\*\*\**p* < 0.0001, \*\*\**p* < 0.001, \*\**p* < 0.01, \**p* < 0.05.

## Supporting Information

Supporting Information is available from the Wiley Online Library or from the author.

## Acknowledgements

Corresponding authors DHR and SR contributed equally to this work. We are grateful to Professor Morag Park (McGill University) for providing us with the cancer cell line used in this study. We gratefully acknowledge the support of the New Frontiers in Research Fund (NFRFE-2020-00470), the Apogee Canada First Research Excellence Fund (CFREF), the Natural Sciences and Engineering Research Council of Canada (NSERC) Discovery Grant (RGPIN-2022-04233), the Canada Summer Jobs program, Fonds de Recherche du Québec – Nature et Technologies (FRQNT) and the Optimizing Power Skills in Interdisciplinary, Diverse & Innovative Academic Networks (OPSIDIAN) studentship funded by the NSERC. We also extend our gratitude to the McGill Scoliosis and Spine Group for their continued support. The NFRF grant was awarded to SR, DHR, and MHW, the Apogee CFREF to SR, the NSERC Discovery grant to DHR, the Canada Summer Jobs program to DHR, the FRQNT scholarship to JBB, and the OPSIDIAN scholarship to LMB. We would like to express our sincere appreciation to Olivia Noik, Pragga Datta, and Markus J. Weber for their assistance with counting red cells in 2D culture using ImageJ for live/dead imaging.

## Data Availability Statement

The data that support the findings of this study are available from the corresponding author upon reasonable request.

