## Supporting Information for "Cold Atmospheric Plasma Selectively Disrupts Breast Cancer Growth in a Bioprinted 3D Tumor Microenvironment Model"

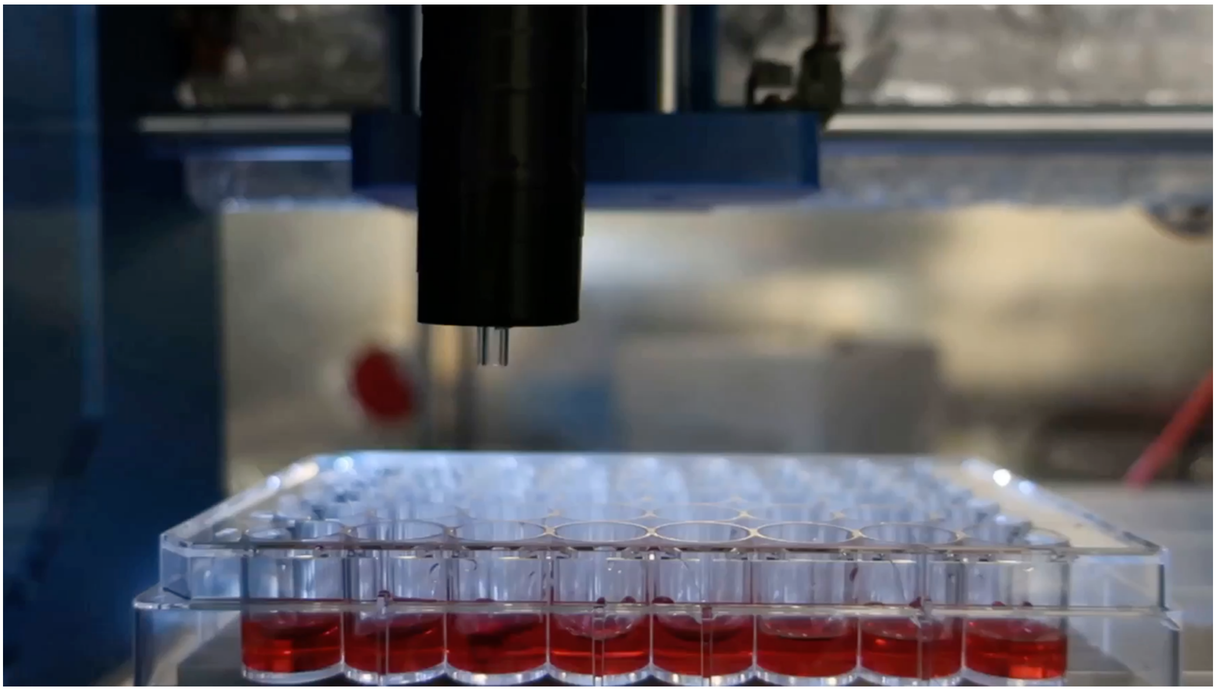

**Figure S1**. Video demonstrating the plasma setup controlled by an XYZ-moving stage for automated treatment of a 48-well plate containing cells.

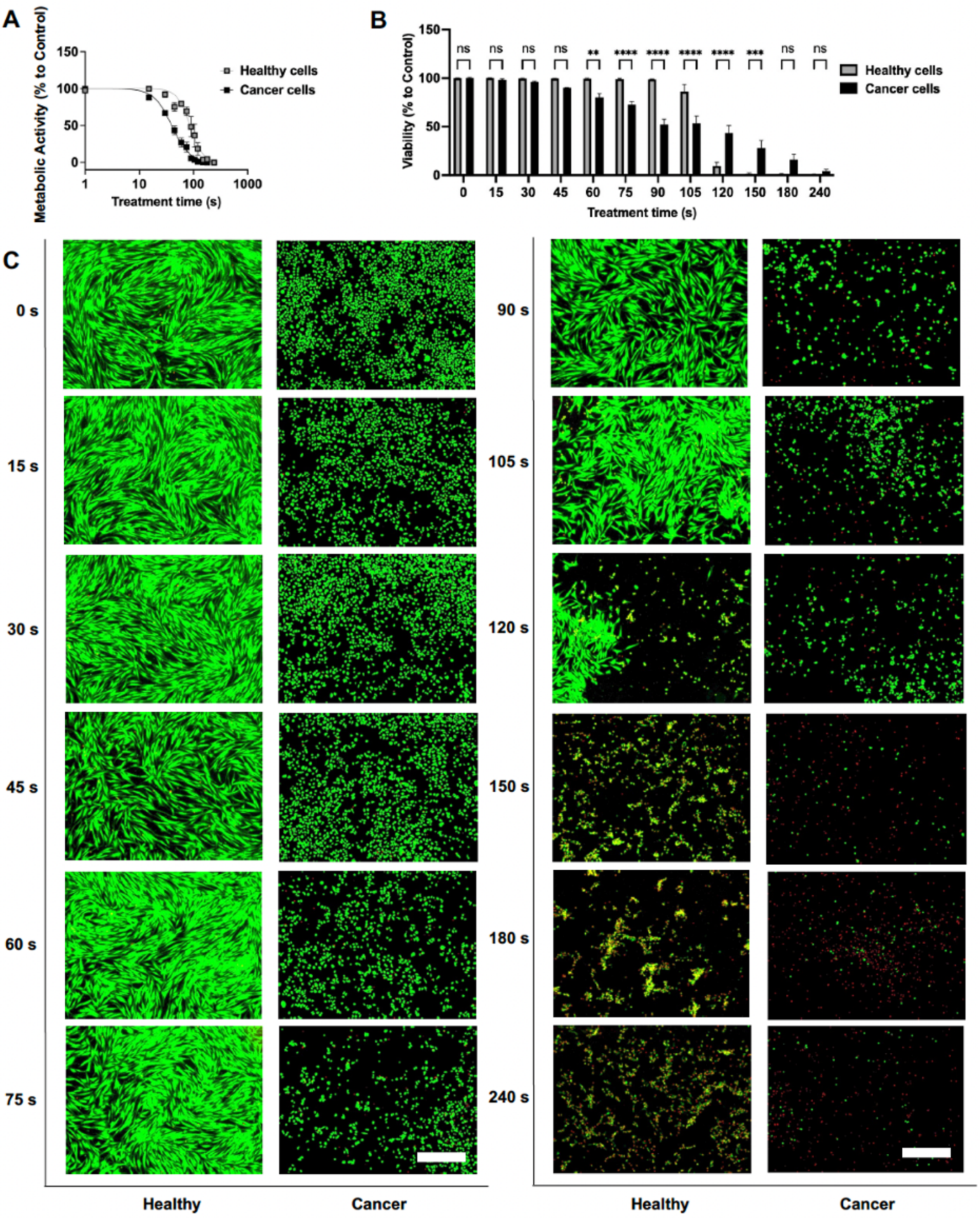


**Figure S2**. Time-dependent effects of CAP treatment on metabolic activity and viability of 2D cultured cancer and healthy cells two days after treatment. A) Time response of the metabolic activity (% compared to control) of cancer and healthy cells for 2 days after plasma treatment of different durations. B) Viability (% compared to control) of cancer and healthy cells for 2 days after plasma treatment of different durations. C) Live/Dead microscopic images (Green: Calcein-AM indicating live cells; Red: Ethidium homodimer-1 indicating dead cells) of the viability of the related cancer and healthy cells from (B), with a scale bar of

750 μm. Statistical significance was determined using two-way ANOVA followed by the Šidák multiple comparison test. Error bars are represented as SEM. Statistical significance is indicated as follows: **** $p < 0.0001$, *** $p < 0.001$, ** $p < 0.01$, * $p < 0.05$; $n = 3$.

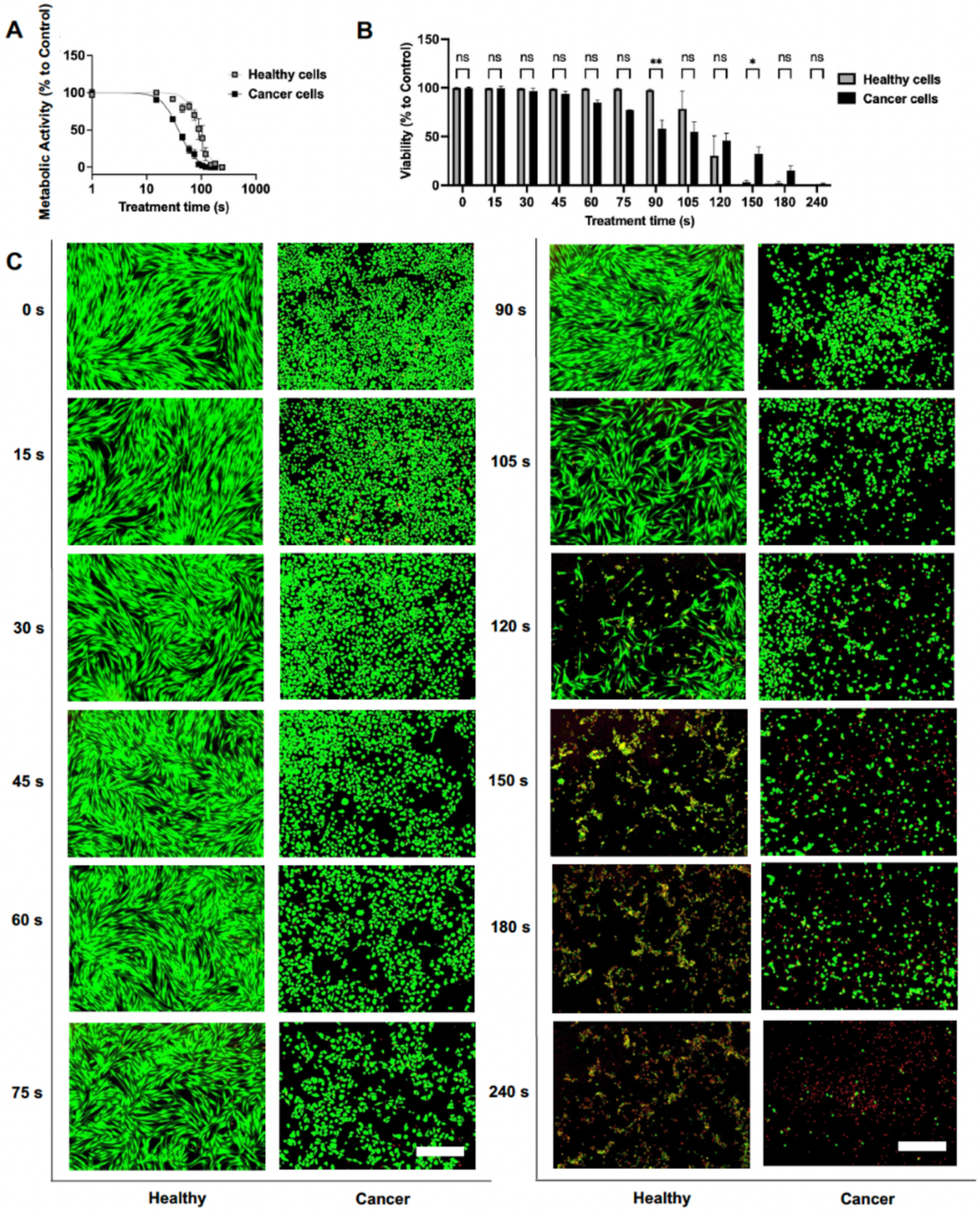


**Figure S3**. Time-dependent effects of CAP treatment on metabolic activity and viability of 2D cultured cancer and healthy cells three days after treatment. A) Time response of the metabolic activity (% compared to control) of cancer and healthy cells for 3 days after plasma treatment of different durations. B) Viability (% compared to control) of cancer and healthy cells for 3 days after plasma treatment of different durations. C) Live/Dead microscopic images (Green: Calcein-AM indicating live cells; Red: Ethidium homodimer-1 indicating dead cells) of the viability of the related cancer and healthy cells from (B), with a scale bar of 750 μm. Statistical significance was determined using two-way ANOVA followed by the Šidák multiple comparison test. Error bars are represented as SEM. Statistical significance is indicated as follows: **** $p < 0.0001$, *** $p < 0.001$, ** $p < 0.01$, * $p < 0.05$; $n = 3$.

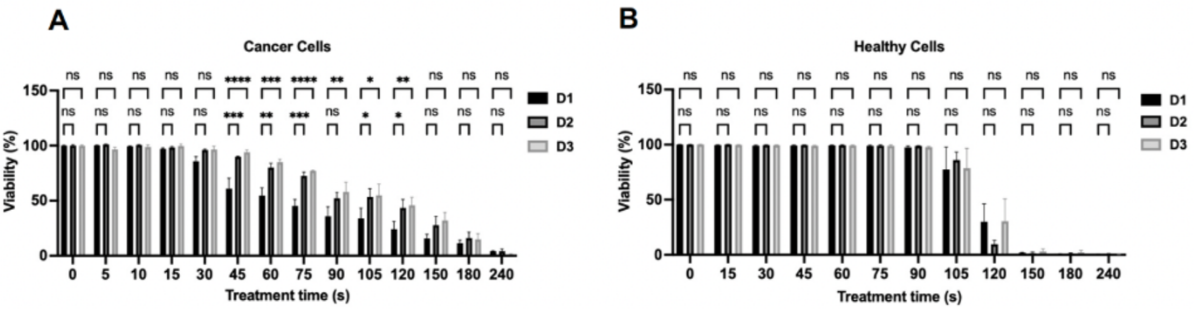


**Figure S4**. Effect of CAP treatment on the viability of 2D cultured cancer and healthy cells over time. A) Viability (% compared to control) of cancer cells for 1, 2, and 3 days after plasma treatment of different durations. B) Viability (% compared to control) of healthy cells for 1, 2, and 3 days after plasma treatment of different durations. Statistical significance was determined using two-way ANOVA followed by the Šidák multiple comparison test. Error bars are represented as SEM. Statistical significance is indicated as follows: **** $p < 0.0001$, *** $p < 0.001$, ** $p < 0.01$, * $p < 0.05$; $n = 3$.

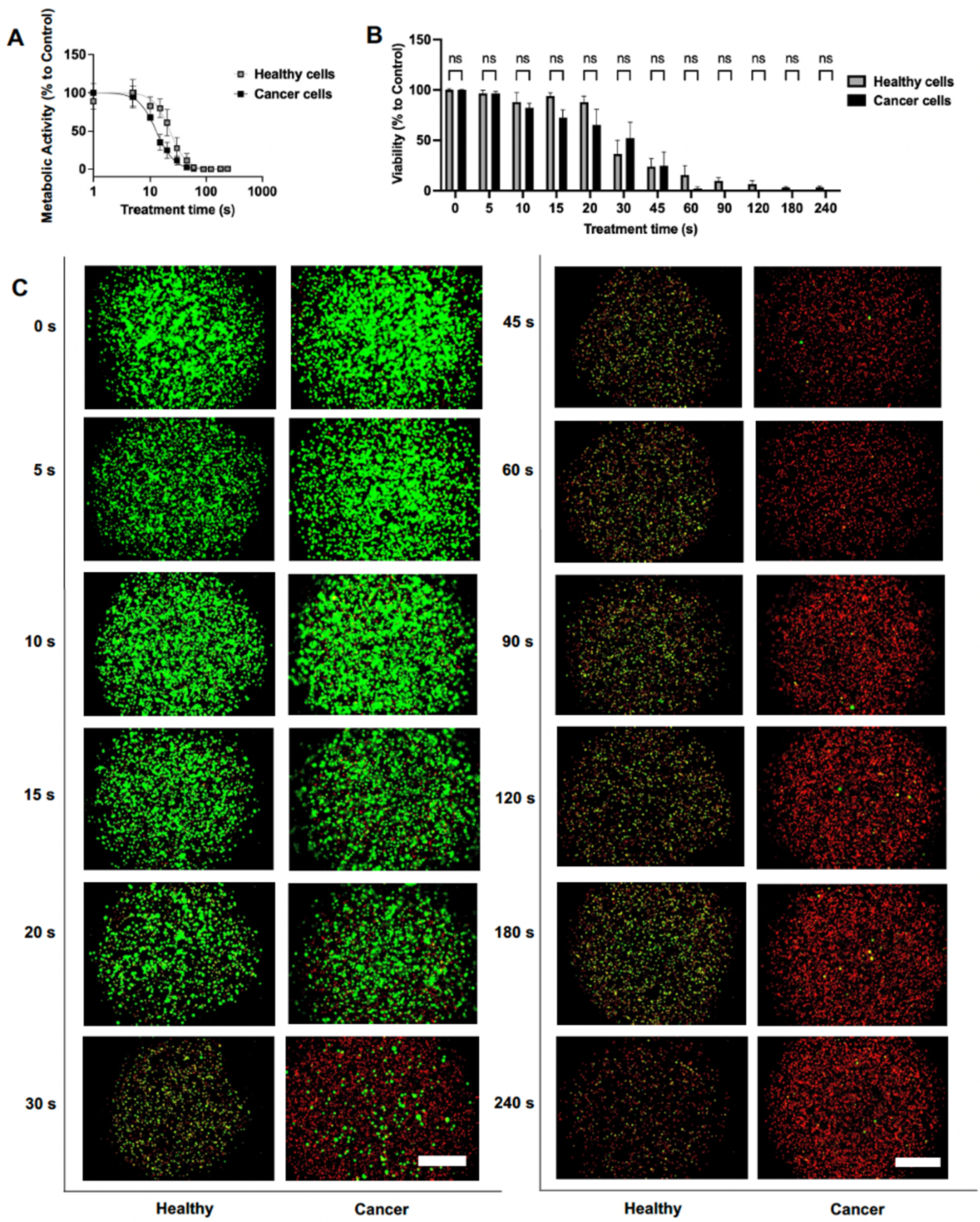


**Figure S5**. Time-dependent effects of CAP treatment on metabolic activity and viability of 3D monocultures of cancer and healthy cells two days after treatment. A) Time response of the metabolic activity (% compared to control) of cancer and healthy cells for 2 days after plasma treatment of different durations. B) Viability (% compared to control) of cancer and healthy cells for 2 days after plasma treatment of different durations. C) Live/Dead microscopic images (Green: Calcein-AM indicating live cells; Red: Ethidium homodimer-1 indicating dead cells) of the viability of the related cancer and healthy cells from (B), with a scale bar of 750 μm. Statistical significance was determined using two-way ANOVA followed by the Šidák multiple comparison test. Error bars are represented as SEM. Statistical significance is indicated as follows: **** $p < 0.0001$, *** $p < 0.001$, ** $p < 0.01$, * $p < 0.05$; $n = 3$.

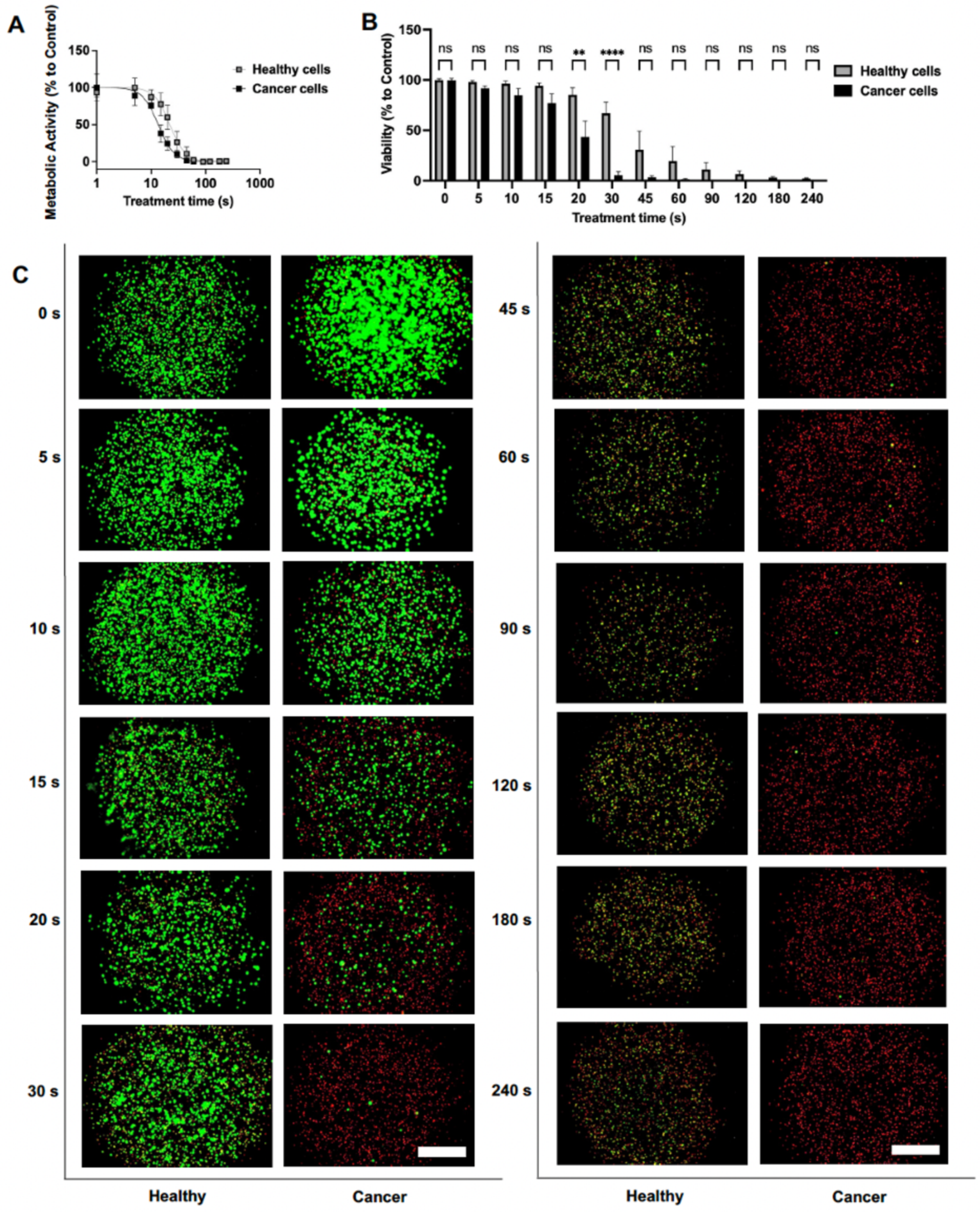


**Figure S6**. Time-dependent effects of CAP treatment on metabolic activity and viability of 3D monocultures of cancer and healthy cells three days after treatment. A) Time response of the metabolic activity (% compared to control) of cancer and healthy cells for 3 days after plasma treatment of different durations. B) Viability (% compared to control) of cancer and healthy cells for 3 days after plasma treatment of different durations. C) Live/Dead microscopic images (Green: Calcein-AM indicating live cells; Red: Ethidium homodimer-1 indicating dead cells) of the viability of the related cancer and healthy cells from (B), with a scale bar of 750 µm, compared to doxorubicin treatment (5 µM). Statistical significance was determined using two-way ANOVA followed by the Šidák multiple comparison test. Error bars are represented as SEM. Statistical significance is indicated as follows: **** $p < 0.0001$, *** $p < 0.001$, ** $p < 0.01$, * $p < 0.05$; $n = 3$.

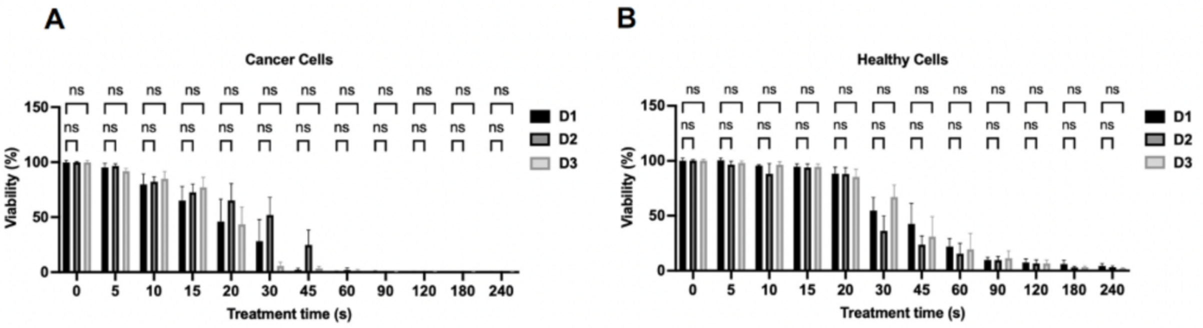


**Figure S7**. Effect of CAP treatment on the viability of 3D monocultures of cancer and healthy cells over time. A) Viability (% compared to control) of cancer cells for 1, 2, and 3 days after plasma treatment of different durations. B) Viability (% compared to control) of healthy cells for 1, 2, and 3 days after plasma treatment of different durations. Statistical significance was determined using two-way ANOVA followed by the Šidák multiple comparison test. Error bars are represented as SEM. Statistical significance is indicated as follows: **** p < 0.0001, *** p < 0.001, ** p < 0.01, * p < 0.05; $n = 3$.

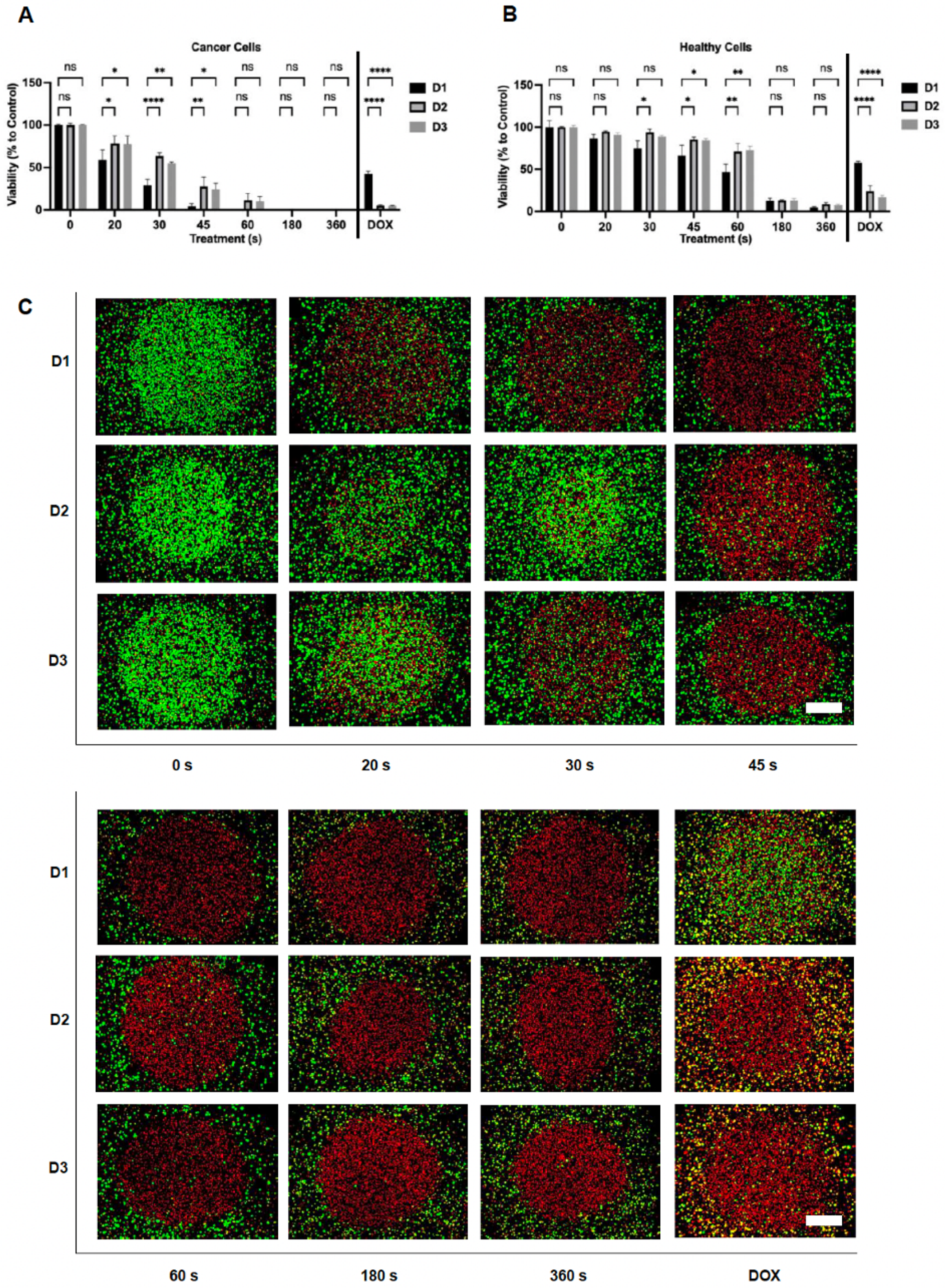


**Figure S8.** Effect of CAP treatment on the viability of 3D co-cultured cancer and healthy cells over time. A) Viability (% compared to control) of cancer cells for 1, 2, and 3 days after plasma treatment of different durations. B) Viability (% compared to control) of healthy cells for 1, 2, and 3 days after plasma treatment of different durations. C) Live/Dead microscopic images (Green: Calcein-AM indicating live cells; Red: Ethidium homodimer-1 indicating dead cells) of cell viability in the co-culture model after 1, 2, and 3 days of CAP treatment at different durations, compared to doxorubicin treatment (5 µM). Scale bar = 1250 µm. Statistical significance was determined using two-way ANOVA followed by the Šidák multiple comparison test. Error bars are represented as SEM. Statistical significance is indicated as follows: **** $p < 0.0001$, *** $p < 0.001$, ** $p < 0.01$, * $p < 0.05$; $n = 3$.

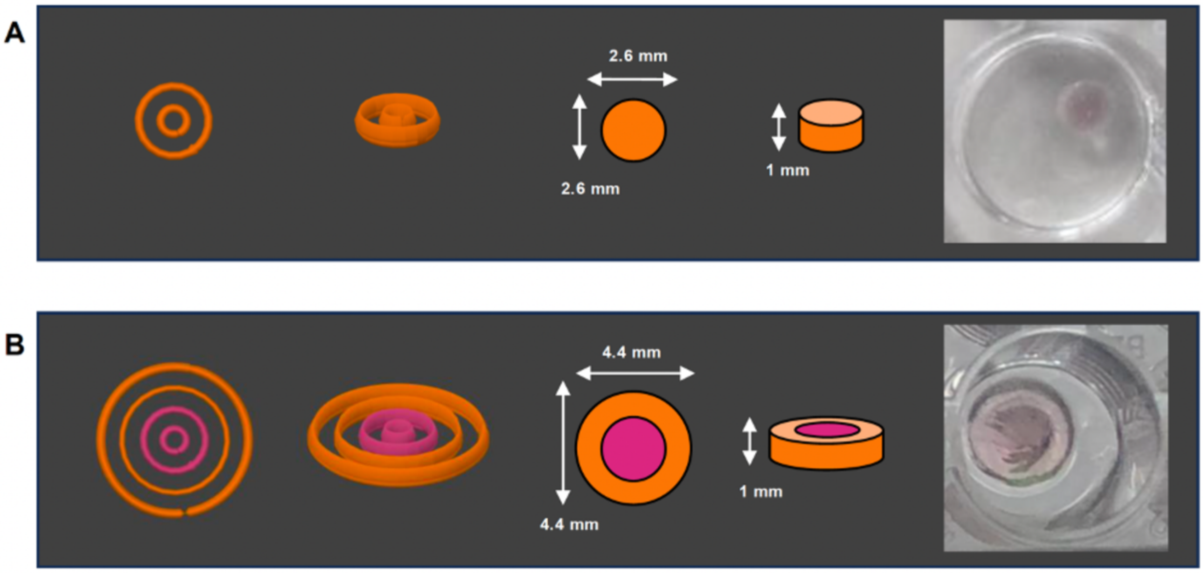


**Figure S9.** Design of the bioprinted models. A) Monoculture model: From left to right, the G-code top view, G-code side view, STL top view, STL side view, and the bioprinted construct in a 48-well plate. B) Co-culture model: From left to right, the G-code top view, G-code side view, STL top view, STL side view, and the bioprinted construct in a 48-well plate. G-codes were visualized using PrusaSlicer, and STL files were visualized with Tinkercad.

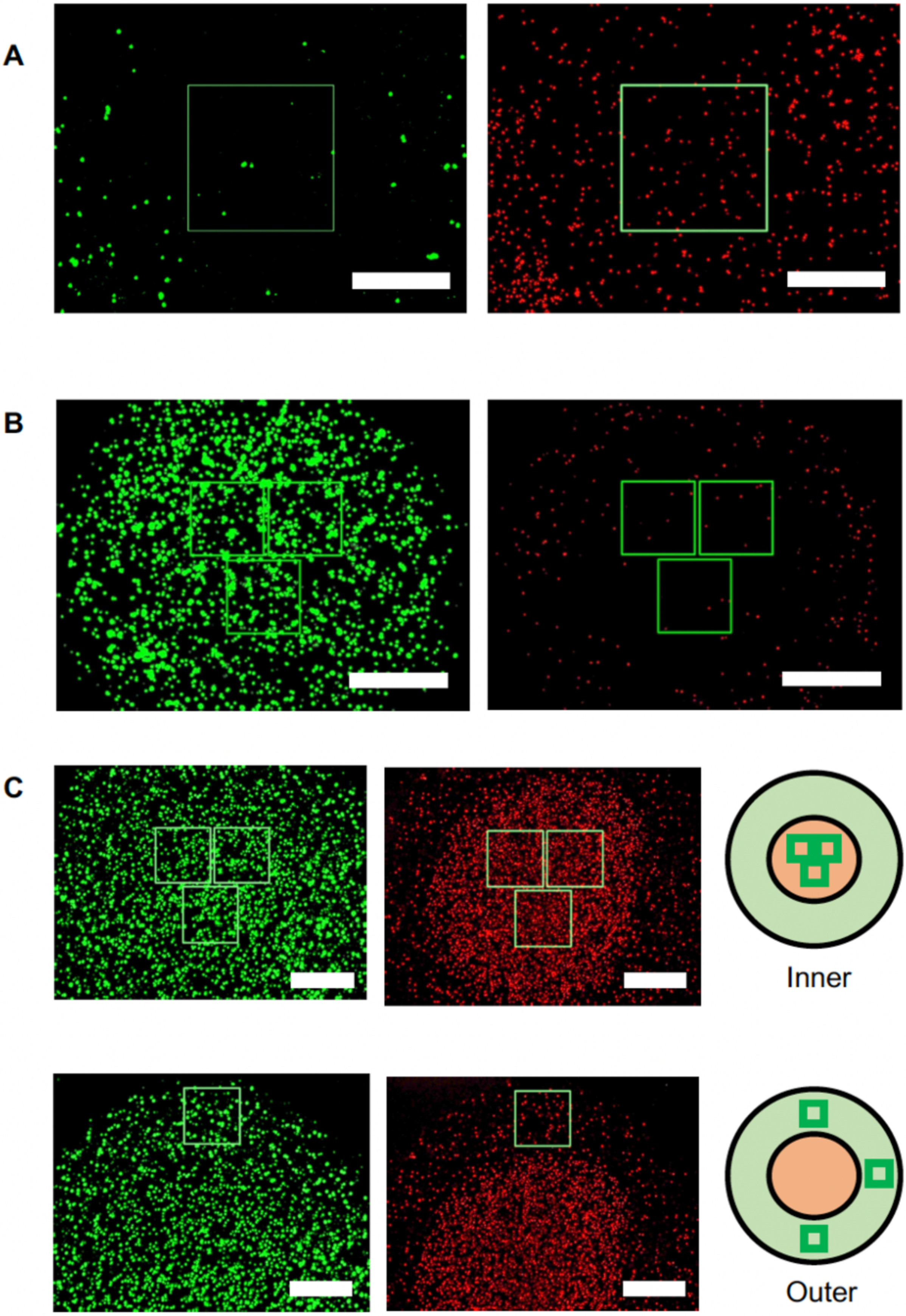


**Figure S10.** Design of programmed squares for cell counting. A) Application in 2D cultures. On the left is the image on the green channel. On the right is the image on the red channel.

Scale bar = 750 µm B) Application in 3D monocultures. Scale bar = 750 µm C) Application in 3D co-cultures. Scale bar = 1250 µm. The Python program draws reproducible squares at consistent locations with specific areas tailored to each culture system, allowing for accurate cell counting within these defined regions.
